# Cryo-EM structure of *ex vivo* fibrils associated with extreme AA amyloidosis prevalence in a cat shelter

**DOI:** 10.1101/2022.05.09.491126

**Authors:** Tim Schulte, Antonio Chaves-Sanjuan, Giulia Mazzini, Valentina Speranzini, Francesca Lavatelli, Filippo Ferri, Carlo Palizzotto, Maria Mazza, Paolo Milani, Mario Nuvolone, Anne-Cathrine Vogt, Giovanni Palladini, Giampaolo Merlini, Martino Bolognesi, Silvia Ferro, Eric Zini, Stefano Ricagno

**Affiliations:** Institute of Molecular and Translational Cardiology, IRCCS Policlinico San Donato, 20097 Milan, Italy; Department of Biosciences, Università degli Studi di Milano, Milan, Italy; Pediatric Research Center Fondazione R.E. Invernizzi and NOLIMITS Center, Università degli Studi di Milano, Milan, Italy; Department of Molecular Medicine, University of Pavia, Pavia, Italy; AniCura Istituto Veterinario Novara, Strada Provinciale 9, 28060, Granozzo con Monticello (NO), Italy; Istituto Zooprofilattico Sperimentale del Piemonte Liguria e Valle d’Aosta, S.C. Diagnostica Specialistica, Via Bologna 148, 10154, Torino, Italy; Amyloidosis Research and Treatment Center, Fondazione IRCCS Policlinico San Matteo, Pavia, Italy; Department for BioMedical Research (DBMR), University of Bern, 3008 Bern, Switzerland; Department of Comparative Biomedicine and Food Sciences, University of Padova, viale dell’Università 16, 35020 Legnaro (PD), Italy; Department of Animal Medicine, Production and Health, University of Padua, viale dell’Università 16, 35020, Legnaro (PD), Italy; Clinic for Small Animal Internal Medicine, Vetsuisse Faculty, University of Zurich, Winterthurerstrasse 260, 8057, Zurich, Switzerland

## Abstract

AA amyloidosis is a systemic disease characterized by deposition of misfolded serum amyloid A protein (SAA) into cross-β amyloid in multiple organs in humans and animals. AA amyloidosis occurs at high SAA serum levels during chronic inflammation. The disease can be transmitted horizontally, likely facilitated by prion-like mechanism, in captive animals leading to extreme disease prevalence, e.g. 70% in captive cheetah and 57-73% in domestic short hair (DSH) cats kept in shelters.

Herein, we present the 3.3 Å cryo-EM structure of an AA amyloid extracted *post-mortem* from the kidney of a DSH cat with renal failure. The structure reveals a cross-β architecture assembled from two 76-residue long proto-filaments. Despite >70% sequence homology to mouse and human SAA, the cat SAA variant adopts a distinct amyloid fold. Based on shared disease profiles and almost identical protein sequences, we propose a similar amyloid fold of deposits identified previously in captive cheetah.

## INTRODUCTION

Amyloidosis is associated with the deposition of proteinaceous amorphous structures in the extracellular space of tissue and organs in humans and animals ^1^. Amyloids in biopsies are histologically revealed by apple-green birefringence under polarized light after Congo Red staining ^1,2^. More than 50 disease-causing amyloidogenic proteins have been discovered, and their molecular identities define specific disease forms and organ distribution ^1,3,4^. Immunohistochemistry and mass spectrometry-based determination of amyloid type is vital for effective treatment ^2,5,6^. The authors of a recent outstanding study have applied single-particle cryo-EM to classify human brain amyloidoses (tauopathies) based on fibril structures, potentially impacting future diagnosis and treatment of these devastating neurodegenerative diseases ^7^. Specifically, AA amyloidosis represents a systemic disease characterized by the deposition of misfolded serum amyloid A protein (SAA) in multiple organs ^2,8^. SAA proteins are 12-14 kDa light apo-lipoproteins that are remarkably conserved throughout vertebrate evolution, indicating critical functions for survival ^9–11^. As part of the host innate response to inflammation, acute-phase variants of SAA (A-SAA) are secreted by the liver to increase serum levels up to 1000-fold ^2,9,11–16^. A minor fraction of A-SAA adopts an α-helical bundle structure that delivers retinol to intestinal myeloid cells, including macrophages, to promote adaptive immunity ^17,18^. The vast majority of A-SAA with a more disordered and enigmatic structure is bound in high-density lipoprotein (HDL), likely contributing to cholesterol homeostasis in macrophages ^2,9,15,19,20^. Chronic inflammation increases A-SAA concentrations to such an extent that macrophages fail to prevent proteolysis-resistant oligomers during lysosomal degradation ^21,22^. Low pH in lysosomes may favor transition of A-SAA into highly ordered almost indestructible amyloid ^21–25^. Final assembly into massive AA amyloid deposits physically distorts and damages organs, in human patients often diagnosed as kidney-related glomerular proteinuria ^2,26,27^. Cryo-electron microscopy (EM) structures of *ex vivo* AA amyloid deposits from diseased organs of a human patient and an experimental mouse model revealed the characteristic cross-β architecture of amyloid, but highly polymorphic structures despite 76% sequence identity ^28^. The two polymorphs were added to a growing amyloid structure database exhibiting more diverse folds than originally anticipated ^25,29^. Proteins of identical sequence may adopt many polymorphs, that are defined *in vitro* by test tube conditions and *ex vivo* by tissue origin and disease type ^3,7,28–32^. Due to the conserved amyloidogenic nature of A-SAA, domestic animals develop systemic amyloidosis similarly to humans ^2,33,34^. Among cats, Siamese and Abyssinian breeds were reported as particularly prone to amyloidosis due to a familial predisposition ^35–40^. Strikingly, the close-to-extinct captive cheetah, from whose lineage DSH cat ancestors split about six million years ago, suffers from an extreme disease prevalence of 70%, likely facilitated by prion-like disease transmission ^41–44^. A prion-like spread of AA amyloidosis was also inferred from studies in which parenteral administered amyloid accelerated deposition in inflamed animals ^2,45,46^. Our recent study has revealed a prevalence of 57-73% among 80 domestic short hair (DSH) cats kept in shelters ^47^, in stark contrast to a very low prevalence (1-2 %) in client-owned cats ^48–51^.

Herein, we present the cryo-EM structure of fibrils extracted *post-mortem* from the diseased kidney of a DSH cat with systemic AA amyloidosis. The structure exhibits the characteristic cross-β architecture of amyloid, but adopts a unique fold distinct from any deposited structure. The novel amyloid fold is built from a SAA variant with potentially increased prion capacity. Almost identical SAA fragment sequences and shared disease profiles hint to a conserved amyloid fold in cat and cheetah.

## RESULTS AND DISCUSSION

### AA amyloid extracted from the kidney of a DSH cat deceased with renal failure

During the last two months of a two-year stay in a shelter in Northern Italy, a female DSH cat became anorectic, developed jaundice and lost significant body weight. She was affected by chronic kidney and liver disease, and had no retroviral infections (Figure S1A). Due to worsened renal failure, euthanasia was requested when the cat was six years old. Histology of the kidney revealed mild chronic multifocal interstitial nephritis and that of the liver showed severe diffuse hypotrophy/atrophy of the hepatocytes. Abundant, amorphous and eosinophilic material in the kidney, liver and spleen stained positive for Congo red and appeared green-apple birefringent under polarized light, consistent with amyloid (Figures 1A and S1B). We suspected AA amyloidosis, representing the most commonly observed type of amyloid in animals ^2,33,34,41,42,45,46^. Indeed, specific antibodies detected SAA close to and as component of amyloids in all three organs (Figures 1B and S1C). Fibrils were extracted from kidney tissue and SAA was identified as the most abundant protein by liquid chromatography with tandem mass spectrometry (LC-MS/MS) (Table S1). Based on negative stain electron microscopy (EM), revealing straight helical filaments with cross-over distances in the 650-700 Å range (Figure 1C), fibril extraction was optimized for collection of a high-resolution single-particle cryo-EM dataset.

**Figure 1.**
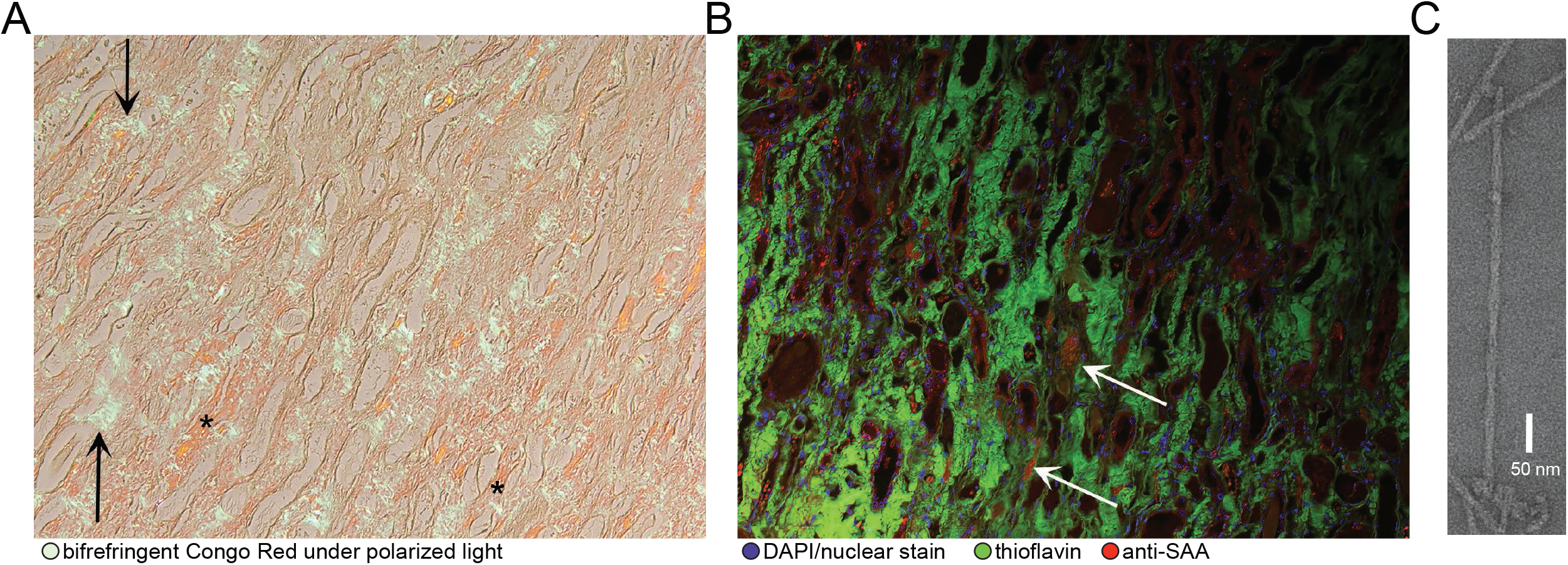
SAA deposits extracted *post-mortem* from the kidney of a shelter cat deceased with renal failure. (A) Abundant interstitial Congo Red-stained amyloid deposits appear orange-red (asterisks) with green-apple birefringence (arrows) under polarized light. Magnification 10x. (B) Immunofluorescent staining of the same kidney slices reveals SAA-positive sections (red) within large areas covered by Thioflavin-stained amyloids (green). Two hotspot areas are highlighted using white arrows. Nuclei are colored blue and were stained using DAPI. Magnification 10x. (C) Micrograph of negative-stained fibril extracted from the kidney. *See related Figure S1 for tissue slices of liver and spleen*.

### Cat’s AA amyloid is built from two identical 76-residue long proto-filaments stabilized through staggered ionic lock and hydrophobic cluster interactions

Cryo-electron micrographs of vitrified AA amyloid extracts revealed a homogeneous population of straight fibrils (Figure 2A) that were manually picked for standard helical reconstruction ^52,53^. About 65k from initially 380k segments were refined with C2 symmetry, a left-handed twist angle of 1.3° and a helical rise of 4.9 Å to yield a final map with a nominal resolution of 3.3 Å, as estimated from half-map Fourier shell correlation curves (FSC) (Figure S2A). Reasonable map-model statistics as well as matching 2D class averages and map projections provide evidence of a physically valid model built into a consistently reconstructed map (Figures 2A-D, S2 and Table S2). The fibril structure is composed of two identical proto-filaments, and exhibits the cross-β architecture characteristic of amyloid (Figure 2). The polypeptide of each proto-filament comprises 11 β-strands between residue positions 19 and 94 and adopts an extended hairpin structure. A central β-arch between residues Asp-50 and Arg-64 links two ∼25 residue long meandering tails that stick together via side chain contacts. A noteworthy feature following the β-arch is an unusual backbone bulge adopted by the P_66_GGAW_70_ segment comprising Pro-66 modeled as *cis*-isomer (Figures S3A), in contrast to the *trans*-Proline residues in mouse and human AA amyloid (Figure S3B). To the best of our knowledge, this is the first example of a *cis*-Proline in amyloid. Isomerization of unfolded SAA may occur spontaneously, as observed in human dialysis-related amyloidosis of β2-microglobulin, but could also be catalyzed by isomerases ^54–59^. In the assembled fibril, the N-terminal tails are surface-exposed at the edges, while the C-terminal tails are buried facing each other (Figure 2C). Each polypeptide deviates from planarity traversing more than three rung layers (Figure 2E). While the β-arch lies almost perpendicular to the fibril axis, the exposed edge- and buried face-tails are tilted by 15° and 10°, respectively. At the intra-protomer interface (Figures 3 and S4, left), the edge-tail of rung layer (i) contacts the face-tails *(i-1)* to *(i+2)*, creating four hydrophobic clusters, three ionic locks and additional H-bond interactions. On the other side, at the inter-protomer interface (Figures 3 and S4, right), the face-tail (i) contacts four rung layers of the adjacent proto-filament, creating two hydrophobic clusters, four ionic locks and two additional H-bond interactions. Such staggered interactions contribute to fibril stability, as described previously ^29^.

**Figure 2.**
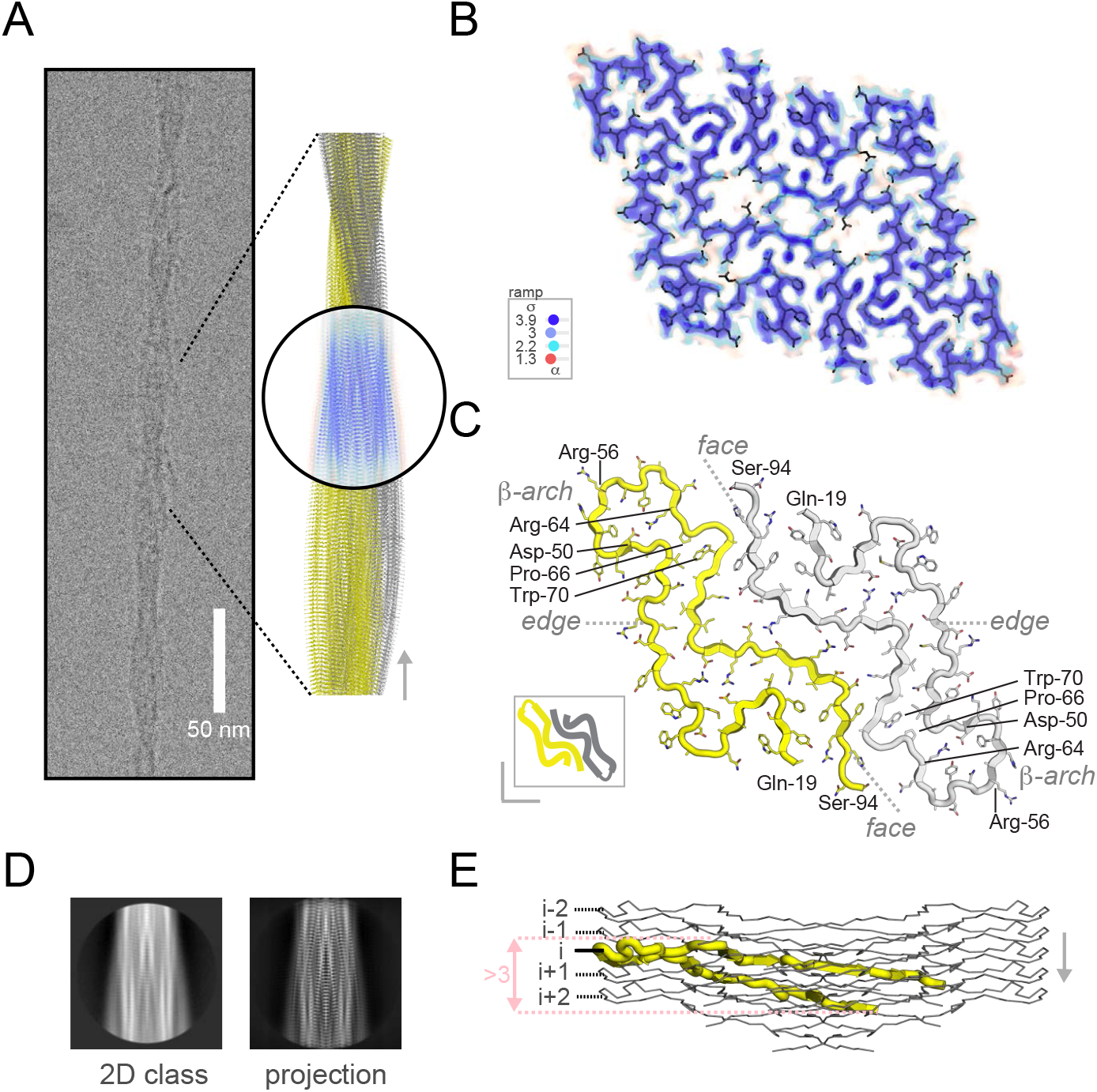
The 3.3 Å resolution cryo-EM structure of the cat’s SAA fibril. (A) Cryo-EM image of a single straight fibril with a crossover distance in the 650-700 Å range. The fibril model spans approximately an entire crossover length of 680 Å and was constructed using the deposited model (PDB: 7ZH7) composed of two proto-filaments (yellow and grey), each assembled by five chains. The map view was oriented to match the fibril orientation of the averaged 2D class and corresponding 2D projection of the reconstructed map. (B) Cross-sectional view of the map volume with contour levels according to the depicted σ-color scale. (C) The molecular model of two subunits within a single fibril layer is shown as cartoon with side chains in yellow and grey. N- and C-terminal positions of each chain and of the β-arch structure are indicated. A scheme in the lower left corner depicts the two chains in yellow and grey. (D) 2D class average corresponding to the orientation of the map shown in panel A. (E) Side-view of the deposited model comprising five subunits in each protofilament. The N- and C-terminal tails are tilted by 10° and 15°, respectively, to the central β-arch that lies almost perpendicular to the long axis of the fibril. Cα-positions of Arg-56 were defined as rung levels (i, i±1 and i±2) along the long fibril axis. *See related Figures S2 and S3 for additional 2D classes, projections, map views, quality indicators and cis-Proline*

**Figure 3.**
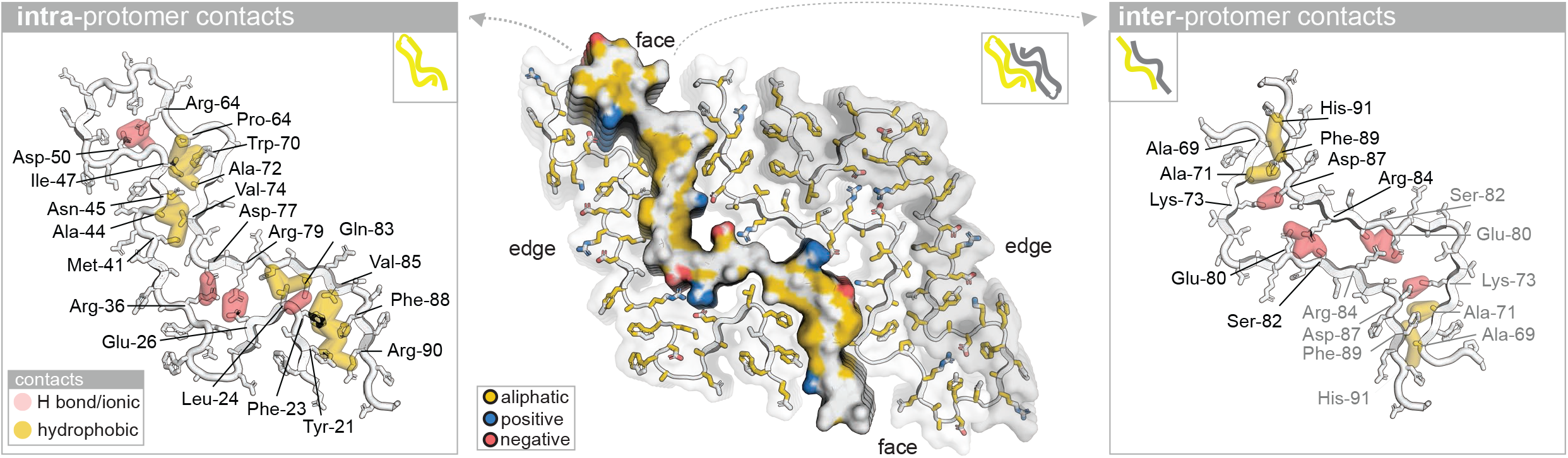
Staggered ionic locks and hydrophobic clusters stabilize intra- and inter-protomer interfaces. (Center) In the cross-sectional view the face of the left proto-filament is represented as molecular surface with aliphatic, positively and negatively charged side chain atoms in yellow, blue, and red, respectively. (Left, right) Side chain contacts at the intra- and inter-protomer interfaces are visualized in separate panels to the left and right, respectively. Hydrophobic and H bond as well as ionic contacts are shown as yellow and pink semi-transparent heavy lines. The backbone and side chain atoms of the opposing strands are represented in mixed cartoon/stick format in white with black outlines. *see related Figure S4 for molecular footprints to illustrate staggered contact modes*

### Cat’s distinct AA amyloid structure buries its unique eight-residue insert between the two proto-filaments and is predicted as the most stable assembly

Although the amino acid sequences of the human, mouse and cat SAA fibrils share >70% residue identity, each amyloid fold is distinct (Figure 4). All three fibrils start at residue 19, but they differ in lengths. Compared to the 54-residue short fibril core of human SAA (hSAA), mouse and cat SAA fibrils (mSAA and cSAA) are elongated by 14 and 22 residues, respectively. Each structure adopts a unique fold, exhibiting distinct arrangements of β-strands that vary slightly in number and lengths, despite high sequence identities (Figure 4A). In each fibril, different parts of the sequences are exposed or buried. In cSAA, residues 19-49 comprising strands β1-β4 are exposed as part of the edge-tail, comprising two short segments that are partially buried in sharp turns. Longer buried segments are observed for the corresponding region in both hSAA and mSAA, but with different distributions. Despite these differences, a segment between residues 24 and 54 of hSAA superposes well on cSAA with an rmsd-value of 2.5 Å (Figure S5). The concomitant observation of shared and distinct structural elements in sequence-homologous amyloids has been referred to as type-2 polymorphism ^29^. The surface-exposed β-arch of cSAA, comprising residues 50 to 64, adopts more extended conformations in hSAA and mSAA. In hSAA, residues 50-55 are buried, followed by the exposed C-terminal segment. In mSAA, residues 50-64 are exposed, while residues 65-86 adopt a U-shaped structure that is, except for residues 66-72, largely exposed and in loose contact with the other protomer. A non-conserved sequence insertion at position 86 of the precursor protein sequence sets apart the cat from mouse and human SAA variants ^10,11^. In the fibril, the insert constitutes a part of the buried tail at the inter-protomer interface. The described differences of the protein sequences, amyloid folds and assemblies yield unique fibril architectures (Figures 4, S6 and S7), each with distinct buried surface area (BSA) and estimated dissociation free energy (ΔG_diss_) contributions of the intra- and inter-protomer interfaces (Figure S8). Compared to mSAA and hSAA, cSAA is predicted as the most stable assembly, with BSA and ΔG_diss_–values increased by ∼2000 Å^2^ and 4-8 kcal/mole, respectively. While the eight-residue insert likely increases fibril stability, predicted local conformational changes may affect the stability of the native lipid-free SAA structure (Figure S9) as well as the currently unknown HDL-bound structure, which might contribute to explain the high amyloidosis prevalence in cats.

**Figure 4.**
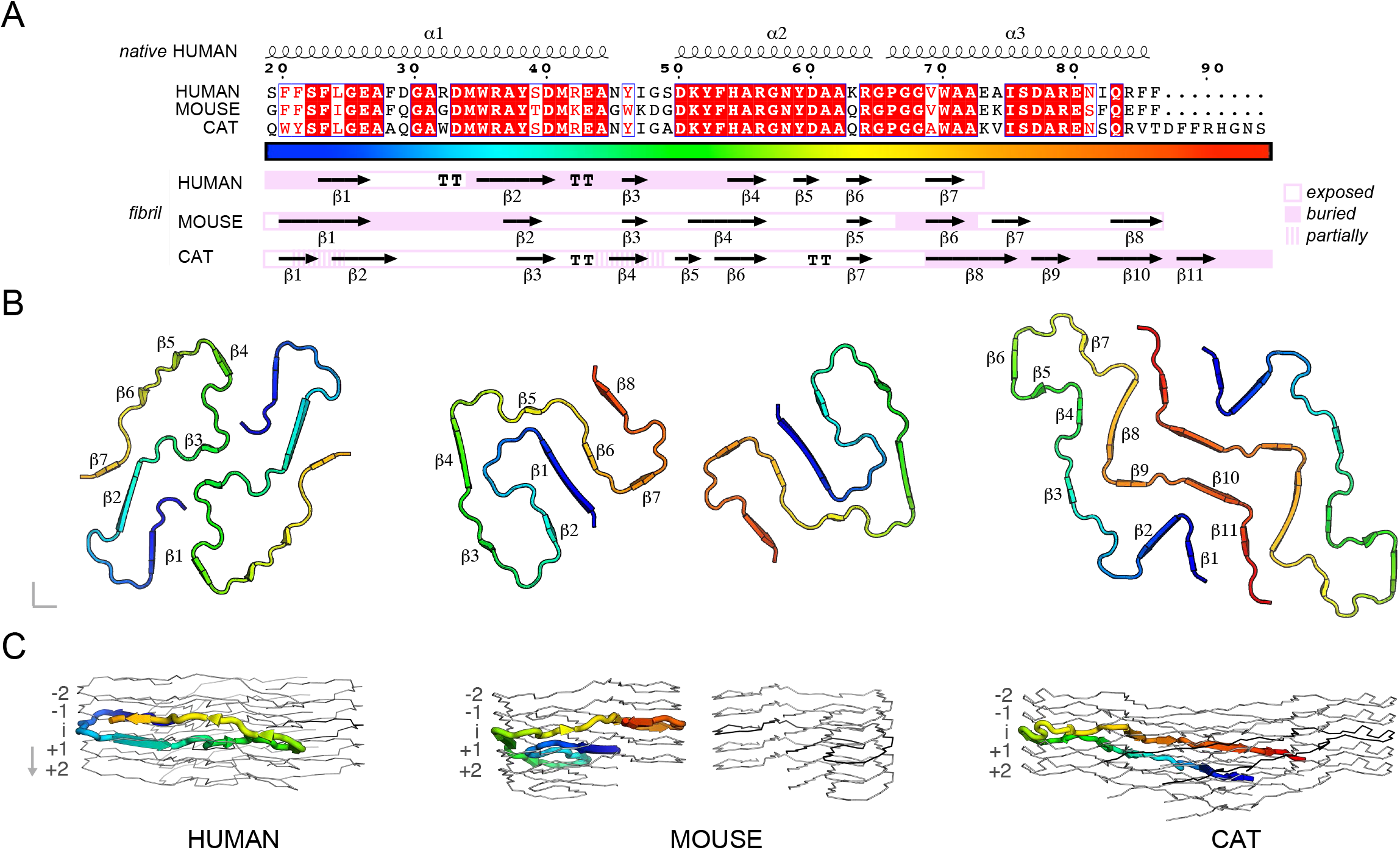
cSAA exhibits weak type-2 polymorphism and buries its unique eight-residue insert in an extended inter-protomer interface. (A) Alignment of hSAA, mSAA and cSAA amino acid sequences present in the fibril core (Uniprot^83^ entries P0DJI8, P05367 and P19707). Strict sequence identity is indicated by a red box with white character, similarities within and across groups are indicated by red characters and blue frames, respectively. For simplicity, numbering is according to cSAA. The alignment was visualized using ESPript^84^. Secondary structure elements of the native human and of the three fibril structures are shown above and below the sequence alignment, respectively. Secondary structures were extracted from PDB^87^ entries 4IP8, 6MST, 6DSO and 7ZH7, respectively. Buried, partially buried and exposed segments were assigned manually taking into account accessible surface areas and relative positioning of segments in the fibril. (B) Cross-section views of human, murine and cat fibrils illustrate the distinct molecular arrangements of strands and interfaces. Residues are colored according to the rainbow code in panel A. (C) Each chain in the human, mouse and cat fibril is not planar, but spans 11, 13.5 and 16.5 Å along the long fibril axis, corresponding to the crossings of about two (human) or three layers (mouse and cat). One chain per fibril is colored as in panel B, the other chains are shown as grey ribbons. *See related Figures S5, S6, S7, S8 and S9 for analysis of shared structural elements, layer level crossing, fibril surfaces, PISA analysis and native SAA structures*.

### Shared disease profiles and almost identical fibril sequences suggest a similar amyloid fold with increased prion capacity in captive cat and cheetah

The presented cryo-EM fibril structure is unique in representing the first *ex vivo* structure of a spontaneously occurring amyloid obtained from an animal kept in a man-made habitat. Remarkably, the distantly related captive cheetah species *Acinonyx jubatus* suffers from a similarly high AA-amyloidosis prevalence of 70%, likely facilitated by a prion-like disease transmission ^41,42,46^. In particular, the amino acid sequence of AA amyloid extracted *post-mortem* from the diseased liver of a cheetah is 97% identical to the sequence of the extracted cat fibril (Figure 5). While highly homologous amyloidogenic proteins, even of identical sequence, may adopt different structure, human brain diseases can be linked to shared amyloid folds ^7,29,30,32^. Based on simple structural considerations we consider the Q19E and N93S substitutions in cheetah fully compatible with the herein presented structure. Although other SAA variants exist in both cat and cheetah, re-discovery of a prion-reported SAA from cheetah in cats affected by severe AA amyloidosis may provide further evidence for its increased prion capacity. Indeed captive cheetah and shelter cats experience similar living conditions that favour horizontal disease transmission, likely through faeces or other exchange of biological material between individuals. Thus, we may hypothesize that the cat and cheetah SAA variant has increased prion capacity with a similar amyloid fold, revealing itself in shelter and zoo populations.

**Figure 5.**
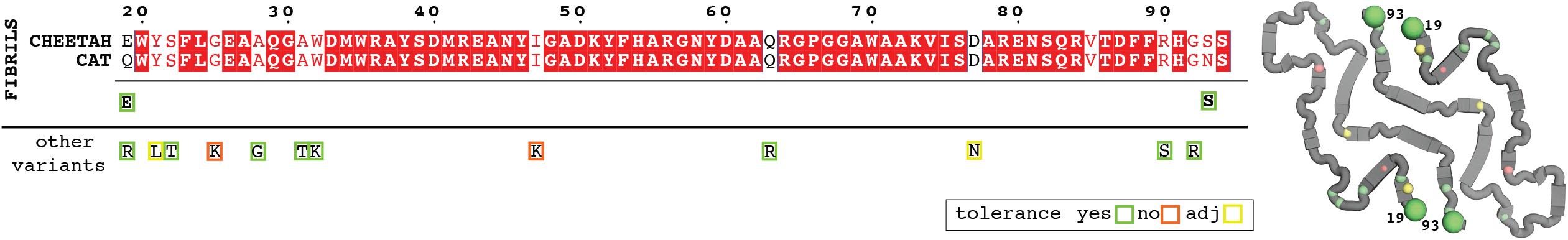
Cheetah AA amyloid fragment is 97% identical to cat fibril. Sequence alignment of extracted cat and cheetah amyloid (Uniprot P19707 and B0M1H2) identified in this and a previous study ^41^. Sequence conservation is based on a multiple sequence alignment ^83,85^ comprising the two AA amyloid as well as eight additional cat and cheetah SAA variants (with Uniprot ids A0A2I2UCY9, A0A6J2AHC5, A0A337S9A8, A0A337SUS3, A0A6J2AJW0, M3WHE0, A0A5F5XYT5, A0A337SKP2). Single-residue substitutions of SAA variants are highlighted on sequence (left) and structure level (right) for fibrils and other reported variants. Substitution tolerance was estimated based on simple structural considerations, and coloured in green, yellow and red. cSAA is shown as grey cartoon, large and small spheres highlight the positions of single-residue substitutions in fibrils and in other cat as well as cheetah SAA variants, respectively.

In summary, here we report the 3.3 Å resolution cryo-EM structure of fibrils from renal tissue of a cat affected by severe AA amyloidosis in a shelter. The fibril is assembled from two twisted proto-filaments, each comprising 76 residues. Amyloid fold and fibril assembly differ from previously reported human and mouse *ex vivo* AA amyloid structures. Almost identical fibril sequences and similar disease prevalence in related captive cheetah suggest that the structure reported here may depict the prion agent responsible for the high AA amyloidosis prevalence in these two related felids.

## MATERIALS AND METHODS

### Diagnosis of AA amyloidosis

#### Histology and immunofluorescence

Full details were described previously ^47^. In brief, organs were collected within 5 h from death, fixed in 10% formalin, and embedded in paraffin. After automatic sectioning, 4-5 μm-thick slices were stained with hematoxylin/eosin and Congo red and examined using standard and polarized light microscopy. For immunofluorescence, serum was obtained by immunization of Balb/c mice with virus-like particles-conjugated to SAA-derived peptides (MREANYIGAD, QRGPGGAWAAKV and EWGRSGKDPNHFRP). Serum specificity was assessed using ELISA. Goat anti-mouse monoclonal IgG conjugated to biotin and streptavidin conjugated to Alexa-546 were used for detection.

#### Fibril extraction

After excision, non-fixed cat kidneys were stored frozen (−80 °C) until amyloids were extracted as described previously ^60–62^. Briefly, 0.5 g tissue from the kidney pole was minced with a scalpel, and washed in 20 mM Tris, 140 mM NaCl, 2 mM CaCl_2_, pH 8. After collagenase-digestion (from *Clostridium histolyticum*, Sigma Aldrich, Saint Louis, MO, USA), the sample was homogenized applying nine cycles of centrifugation and pellet re-suspension in 1 mL of 20 mM Tris, 140 mM NaCl, 10 mM EDTA, pH 8.0. Supernatants from additional homogenization cycles in ice-cold water were kept as amyloid extracts and analyzed by SDS-PAGE.

#### LC-MS/MS

Extracted fibrils were solubilized in 8M Urea, 0.1M Dithiotreitol and quantified using Bradford (Bio-Rad, Hercules, CA, USA). 30 μg of solubilized and reduced protein was alkylated (150 mM iodoacetamide, 1 h, RT, dark), 1/6-diluted in 100 mM NH_4_HCO_3_, and digested with Trypsin (Sequence grade, Promega, Madison, WI, USA) at a 1:20 (w/w) ratio for 16 h at 37 °C. Peptides were purified using Pierce C18 Tips (Thermo Fisher Scientific) and analyzed by LC-MS/MS (Table S1). Uniprot entries Q9XSG7, Q1T770, A0A337SKP2 and Q5XXU5, were identified as top hits from the *Felis catus* proteome.

### Structure of AA amyloid fibrils by single-particle cryo-EM

#### Sample preparation and data collection

A 4-μl droplet of fibrils sample was applied onto a C-flat thick 1.2/1.3 300 mesh Cu, previously glow-discharged for 30s at 30mA using a GloQube system (Quorum Technologies). The sample was blotted immediately and plunge-frozen in liquid ethane using a Vitrobot Mk IV (Thermo Fischer Scientific). A cryo-EM dataset of 2,652 movies was collected automatically on a Talos Arctica 200kV (Thermo Fisher Scientific), equipped with a Falcon 3 direct electron detector operated in electron counting mode (Table S2).

#### Helical reconstruction

Fibrils were picked manually from dose-weighted, motion- and CTF-corrected image micrographs in RELION 3.1 ^52,53,63,64^. After manual picking, a first set of ∼65,131 segments were extracted in 1000-pixel boxes binned by 4 and a 10% inter-box distance. The tube diameter, rise and number of asymmetrical units were set to 125 Å, 4.75 Å and 21, respectively. Reference-free 2D classification was performed to select a single large class average for initial model generation with an estimated cross-over distance of 700 Å. A second set of ∼381,233 smaller segments was extracted for the refinement applying a box size of 250 pixel with 10% inter-box distance and helical tube diameter, rise and asymmetrical unit values of 150 Å, 4.75 Å and 5, respectively. The initial model was re-scaled and re-windowed to match the un-binned particles and low-pass-filtered to 10 Å. 3D auto-refinement applying C1 symmetry, angular sampling, helical twist and rise values of 3.7°, 1.3° and 4.75 Å, respectively, yielded an ∼4 Å resolution map. Imposing apparent C2 symmetry improved map resolution to 3.8 Å. After additional steps comprising 3D class average selection, Bayesian polishing, CTF refinement and mask-generation, ∼65,122 particles were subjected to a final 3D auto-refinement with solvent-flattened FSCs. The final map was reconstructed with helical twist and rise values of 1.3° and 4.9 Å to an estimated resolution of 3.3 Å.

#### Model building

After map auto-sharpening in Phenix ^65^, the model was built *de novo* starting from a map region featuring an unusual backbone bulge with an associated bulky side-chain volume. The bulge was identified as P_66_GGAW_70_ in the LC-MS/MS-identified amino acid sequence. The model was built and refined in Coot, Chimera-Isolde as well as Phenix real-space refinement initially with and later without Amber gradients ^66–70^. Molprobity validation^71^ revealed model issues that were resolved by rebuilding of a single chain into the inverted map with left-handed twist. Five 76-residue long chains in each proto-filament were modeled and refined with non-crystallographic symmetry (NCS) restraints. In the final stages of refinement, we modeled Proline-66 as *cis*-isomer to fit the backbone carbonyl into the map, although a higher resolution is required to discriminate conclusively between *cis-* and *trans*-Proline. Phenix, Molprobity and EMDB validation^71–73^ revealed map-model cross-correlation (CC_mask_), EM-ringer and Molprobity-score values of 0.74, 5.1 and 1.4, indicative of a physically valid model with definite map support.

### Data analysis and visualization

Structures and derived data were analyzed and visualized using PyMol and Rstudio ^74–79^. Molecular contact fingerprints, flexible structural alignments and buried surface areas as well as dissociation free energies of assemblies were obtained from Arpeggio, FATCAT and PISA webservers ^80–82^. Sequences were aligned and visualized using Uniprot, Blast, ClustalOmega and ESPript ^83–86^.

## ACKNOWLEDGEMENTS

This study was partially supported by Ricerca Corrente funding from Italian Ministry of Health to IRCCS Policlinico San Donato; by Centro di Ricerca Pediatrica, Fondazione Romeo and Enrica Invernizzi (Milan, Italy); Fondazione ARISLA (project TDP-43-STRUCT); Italian Ministry of Research PRIN 2020 (20207XLJB2); AniCura Clinical Research Grant 2021.

