## supplementary figures and tables for "Cryo-EM structure of *ex vivo* fibrils associated with extreme AA amyloidosis prevalence in a cat shelter"

### SUPPLEMENTARY TABLES AND FIGURES

**Table S1. Top hits identified by LC-MS/MS and data acquisition parameters**

| | |
|---|---|
| **Top hits** | |
| Uniprot entries ranked by relative abundance | (1) ***Q9XSG7***, (2) ***Q1T770***, (3) ***A0A337SKP2***, (4) ***Q5XXU5***, |
| **Data acquisition parameters** | |
| **Liquid chromatography (LC)** | |
| UHPLC instrument | Dionex Ultimate 3000 |
| Column | PepMap RSLC C18 |
| (particle, pore sizes, dimensions) | (2 µm, 100 Å, 75 µm x 50 cm) |
| linear gradients | 2-40% formic acid 0.1% in acetonitrile in 70 min a[illegible] from 40-95% in 10 min, |
| flow rate | 250 nL/min. |
| **MS/MS** | |
| Instrument (Company) | Q Exactive (Thermo Fisher Scientific) |
| Data acquisition | full mass spectra in positive ion mode |
| m/z range | 400-1500 |
| resolution | 70000 FWHM |
| automatic gain control | $10^6$ |
| maximum injection time | 120 ms |
| Higher-energy C-trap dissociation (HCD) | |
| fragment selection (charge states) | 10 most intense precursor ions (+2, +3 and +4) |
| normalized collision energy | 27% |
| isolation window | 2 m/z |
| Dynamic exclusion | 30 s |
| Fragment spectra | |
| resolution | 17500 FWHM |
| automatic gain control | $5*10^4$ |
| maximum injection time | 250 ms |
| **Data processing** | |
| Software | Proteome Discoverer software, version 2.0 |
| Proteome (Uniprot) | *Felis catus* (UP000011712) |
| Search settings | Semi-tryptic cleavage |
| Max missed cleavages | 2 |
| Precursor mass tolerance | 10 ppm |
| Fragment mass tolerance | 0.03 Da |
| Static modification | Carbamidomethylation (+57.021) of cysteines |

**Table S2. Cryo-EM data acquisition parameters and map/model statistics**

| | |
|---|---|
| **Data acquisition and processing** | |
| **Data collection and image processing** | |
| Microscope (Company) | TALOS Arctica (Thermo Fischer Scientific) |
| Voltage (kV) | 200 |
| Camera | Falcon 3EC |
| Data collection software | EPU-2.8 |
| Magnification | × 120,000 |
| Total electron dose ($e^-/Å^2$) | 40.0 |
| Defocus range (µm) | -0.8 and -2.2 µm |
| Pixel size (Å) | 0.889 |
| Micrographs (no.) | 2,652 |
| **Data processing and reconstruction** | |
| Software | Relion 3.1 [1,2] |
| dose-weighting and motion-correction | MOTIONCOR2 [3] |
| CTF-correction | CTFFIND4 [4] |
| Helical reconstruction | |
| Imposed symmetry | C2 |
| Twist / rise | 1.3° / 4.9 Å |
| Initial segments (no.) | 381,233 |
| Final segments (no.) | 65,122 |
| Phenix auto-sharpening B-factor ($Å^2$) | 139.07 |
| **Map/model statistics** | |
| **Model refinement** | |
| r.m.s. deviations | |
| Bond lengths (Å) | 0.003 |
| Bond angles (°) | 0.941 |
| Ramachandran plot | |
| Favored (%) | 95.95 |
| Allowed (%) | 4.05 |
| Outliers (%) | 0.00 |
| Validation | |
| Molprobity score | 1.40 |
| Clashscore | 3.28 |
| Poor rotamers (%) | 0.00 |
| EM-Ringer Score | 5.1 |
| Map-model correlation | |
| CC mask/box/peaks/volume | 0.74 / 0.68 / 0.68 / 0.74 |
| **Deposition codes** | |
| PDB code | 7ZH7 |
| EMDB code | EMD-14726 |
| EMPIAR | EMPIAR-11001 |

**Figure S1. Congo Red and immunofluorescent staining of amyloid in spleen and liver**

(A) Red blood cell count, serum bilirubin and creatinine concentrations of the cat and reference range (red).

(B) Congo red and (C) immunofluorescent staining of spleen (top) and liver (bottom) reveal amyloid deposits in both organs. B) Amyloid deposits appear orange-red (black arrows) with green-apple birefringence (white arrows) under polarized light. Magnification 20x. In the liver, the hepatocytes (white arrows) are separated and isolated by abundant amyloid deposits that appear orange-red (asterisk) with green-apple birefringence (black arrows) under polarized light. Magnification 20x.

**Figure S2. 2D class averages, Fourier shell correlation curves and residue-level map-model cross-correlation plots indicate a consistent reconstruction yielding a model with definitive map support**

(A) Half-map (black) and model-map (grey) based Fourier shell correlation (FSC) curves from EMDataBank and Phenix [5,6] yield global resolution estimates of 3.3 Å.

(B) A residue-level map-model cross-correlation plot reveals map support for all residues, with a mean value of 0.73±0.04. Map-model CC values of >0.7 and <0.5 are thought of as good and poor fits, respectively [6].

(C) Images of three additional 2D class averages are compared to the 2D projections and volume representations of the reconstructed map, as well as model visualizations as in Figure 2A. Map orientations were matched by applying the known rotation angles from the 2D projections. Rotation angles to turn the fibril around the long axis, relative to the orientation in Figure 2D, are indicated. Thin red lines were added to the top 2D class average to highlight the non-staggered rung levels. Staggering would be expected for pseudo-$2_1$ symmetry, as described previously [7].

**Figure S3. *cis*-Proline is supported by map and side chain pointing directions of following residues**

(A) As test Pro-66 was modeled both as *trans* and *cis*-isomer with similar Molprobity quality statistics, however the *cis*-isomer appeared as a better fit to the map. Map contours were colored as in Figure 2B.

(B) The side chains of residues following Pro66 in cSAA point to opposite directions compared to the mSAA and hSAA structures, where Pro66 is in *trans* conformation. The side chain pointing directions are shown for hSAA, mSAA and cSAA from top to bottom in two separate panels. (left panel) Residues 62-70 are visually aligned to compare side chain pointing directions. Up- and down directions are defined based on the backbone shown as cartoon. (right panel) Upwards, and downwards pointing side chains are shown in blue and red.

**Figure S4 Footprints of central rungs on the face sheet at the intra- and inter-protomer interfaces**

Molecular footprints illustrate the staggered contacts between the central edge and face rungs at the intra- and inter-protomer interfaces, respectively. The molecular surfaces are colored (left) as in Figure 3, and (right) on a blue-green color-ramp based on the distances to the opposing central rungs.

**Figure S5. Short structural elements shared between cSAA and hSAA**

A single 20-residue long segment from residues 24 to 54 of h SAA is superimposed on cSAA with an rmsd-value of 2.5 Å, and shown as black ribbon.

**Figure S6. Layer level crossings of non-planar rungs in hSAA, mSAA and cSAA**

The rungs of all three fibrils are non-planar and their backbone C$\alpha$-positions cover distance ranges of 11, 13.5 and 16.5 Å along the long axes of the human, mouse and cat fibrils (A) Same view as in Figure 4C, but the chain is shown in black.

(B) In case of the human and mouse fibrils the position of backbone C$\alpha$ atoms of a single chain is plotted along the long fibril axis (red coordinate system). The layer lines are based on the C$\alpha$ positions of Ala-31 and Gly-57. Due to the large tilt angles in the cat fibril, the C$\alpha$-positions of the left protomer are turned by 14° to re-orient the N-terminal tail perpendicular to the viewing y-axis (green coordinate system).

**Figure S7. Molecular polymorphs with distinct intra- and inter-protomer interfaces and fibril architectures**

(A) The distinct intra- and inter-protomer interfaces of each fibril are apparent in cross-section views of single rungs in cartoon/stick format on the yrb-scale. Residues accessible on the surface of assembled fibrils are labeled.

(B) Assembly of the polymorphic human, mouse and cat SAA rungs into fibrils yields structures with distinct surface chemical properties. Surface-accessible residues are labeled as in panel A. Pitch lengths are indicated.

**Figure S8. Larger buried surface area renders cSAA as the thermodynamically most stable fibril**

(A) Cross-sectional and (B) side views for the assembly of single chains (red) within each proto-filament (yellow) and fibril (yellow+grey) visualize the intra- and inter-molecular interaction surfaces within the hSAA, mSAA and cSAA. Bar plots compare buried surface areas (bsa) and dissociation free energies ($\Delta G_{diss}$) of single chains within each proto-filament (yellow) and the additional contributions of the inter-protomer interface (grey). Values were obtained from PISA[8].

**Figure S9. Local conformational changes in the predicted structure of cat´s native lipid-free SAA comprising the eight-residue insert**

(A) Native human SAA adopts a four-helix ($\alpha$1-$\alpha$4) bundle structure (PDB: 4IP9) [9]. The amyloid-forming segment is highlighted in magenta. (B) The AI-based model [10] of native cat SAA differs in the conformations of the start and end segments of helices $\alpha$4 and $\alpha$3, respectively, as well as of the connecting loop, due to the presence of the eight-residue insert shown in pink.

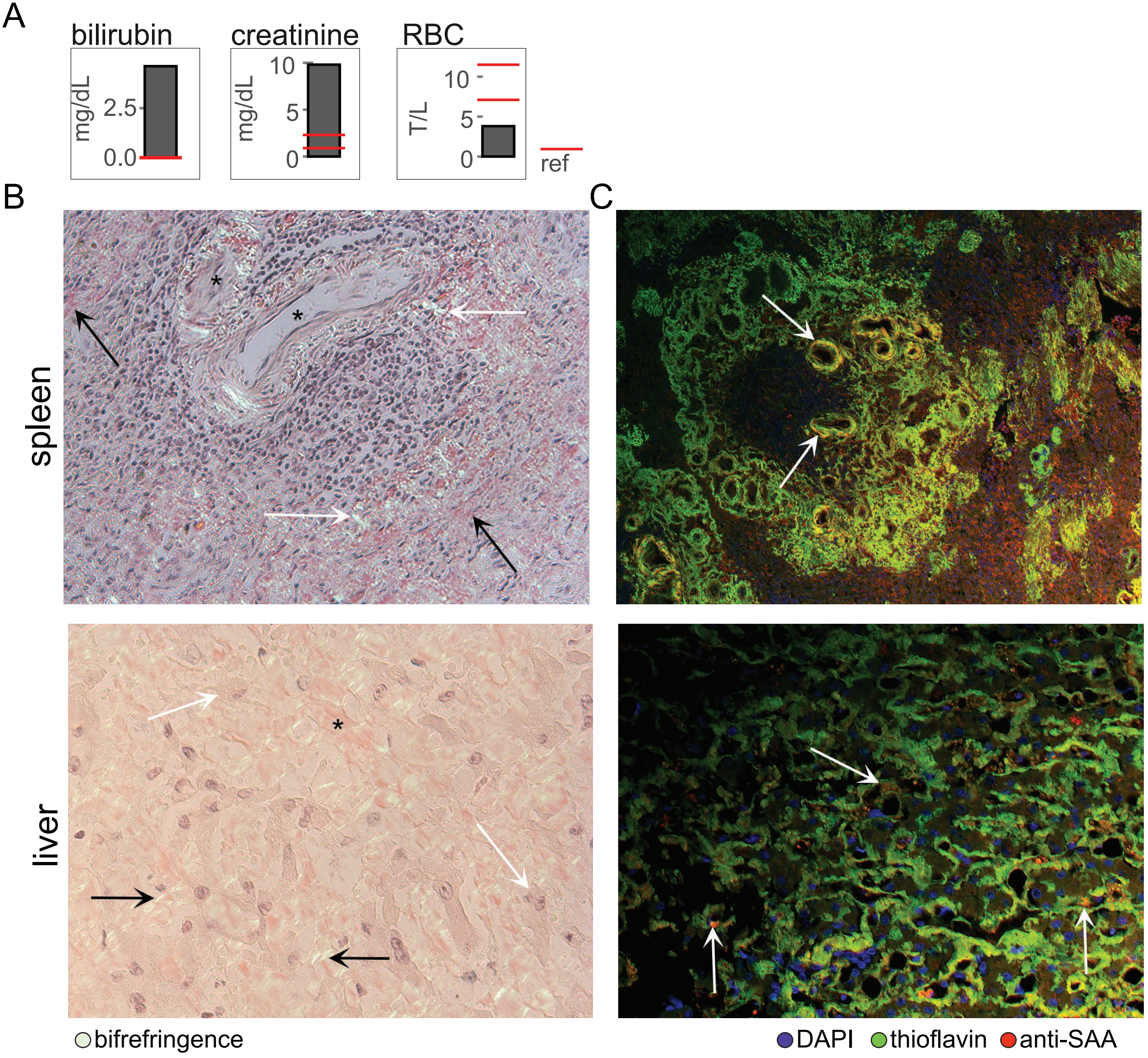
A
bilirubin
mg/dL
2.5
0.0
creatinine
mg/dL
10
5
0
RBC
T/L
10
5
0
ref
B
C
spleen
liver
bifrefringence
DAPI
thioflavin
anti-SAA

Figure S1

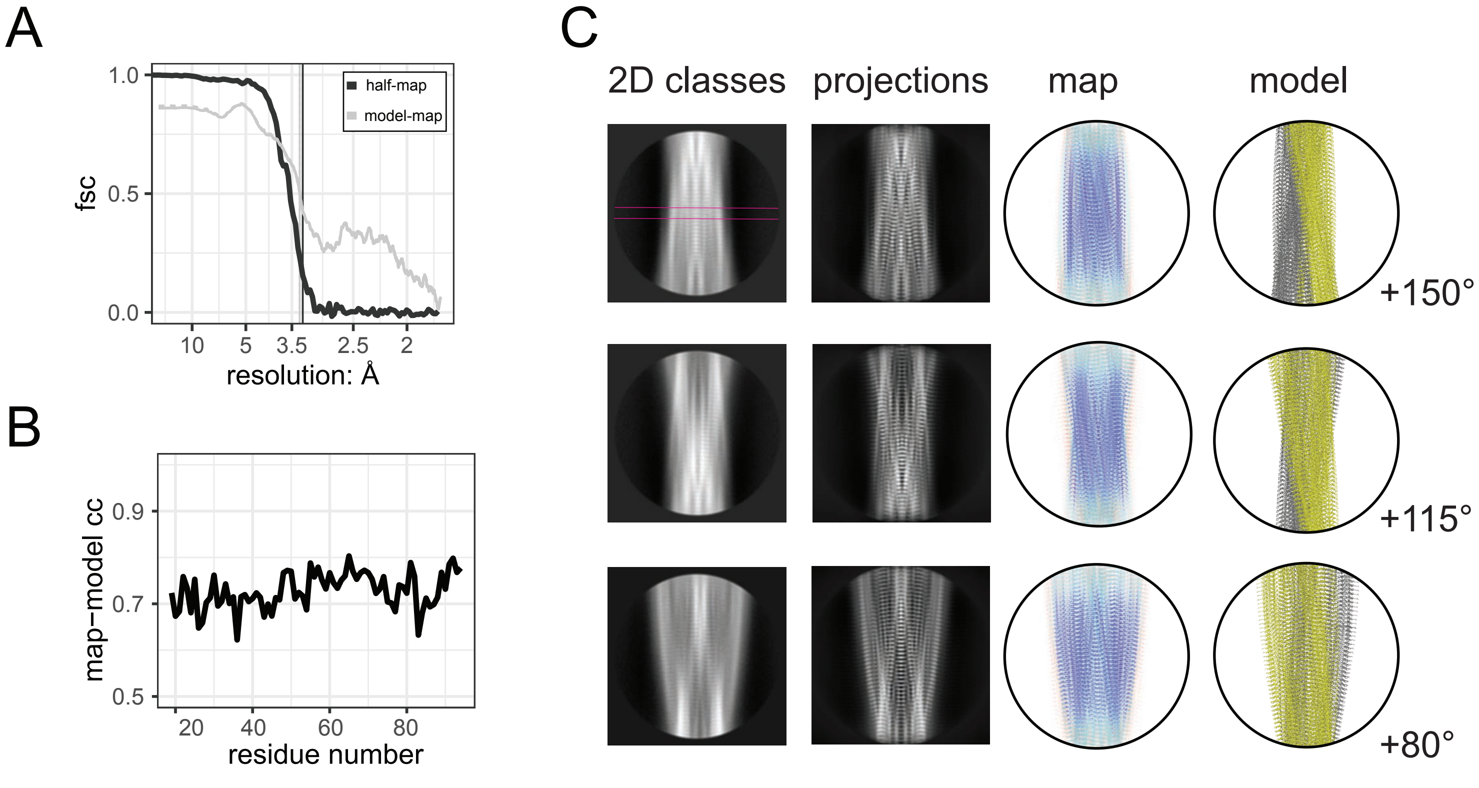
A
1.0
0.5
0.0
fsc
half-map
model-map
10
5
3.5
2.5
2
resolution: Å
B
0.9
0.7
0.5
map−model cc
20
40
60
80
residue number
C
2D classes
projections
map
model
+150°
+115°
+80°

Figure S2

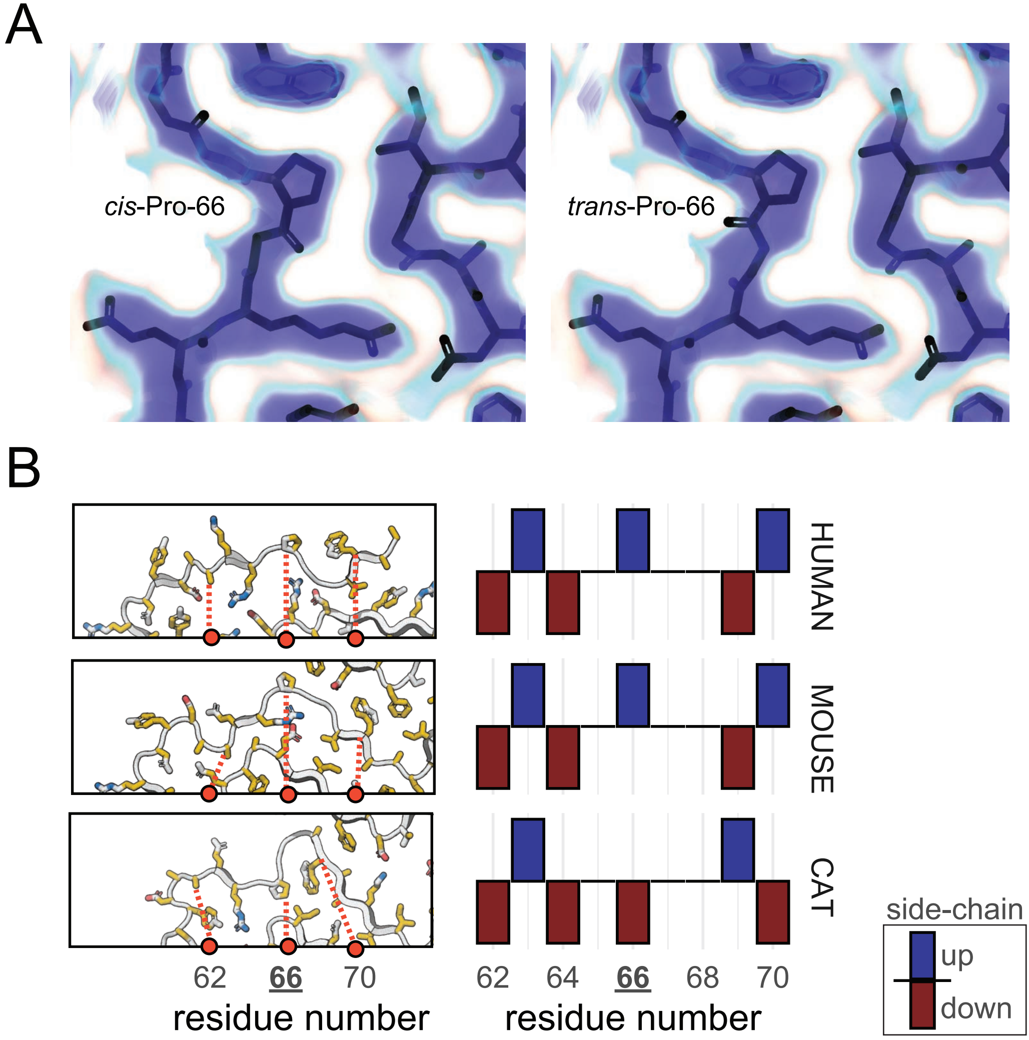
A
cis-Pro-66
trans-Pro-66
B
HUMAN
MOUSE
CAT
62
66
70
residue number
62
64
66
68
70
residue number
side-chain
up
down

Figure S3

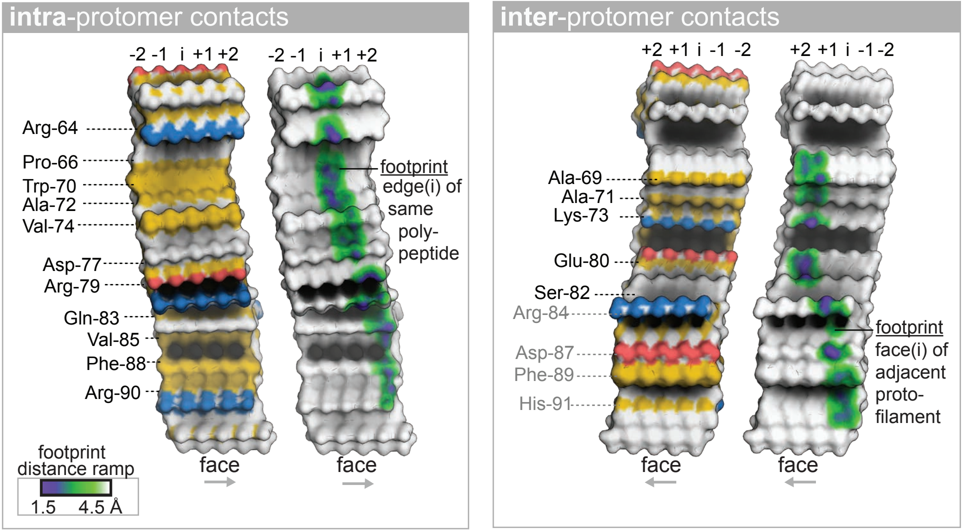
intra-protomer contacts
-2 -1 i +1 +2
-2 -1 i +1 +2
Arg-64
Pro-66
Trp-70
Ala-72
Val-74
Asp-77
Arg-79
Gln-83
Val-85
Phe-88
Arg-90
footprint
edge(i) of
same
poly-
peptide
face
face
footprint
distance ramp
1.5
4.5 Å
inter-protomer contacts
+2 +1 i -1 -2
+2 +1 i -1 -2
Ala-69
Ala-71
Lys-73
Glu-80
Ser-82
Arg-84
Asp-87
Phe-89
His-91
footprint
face(i) of
adjacent
proto-
filament
face
face

Figure S4

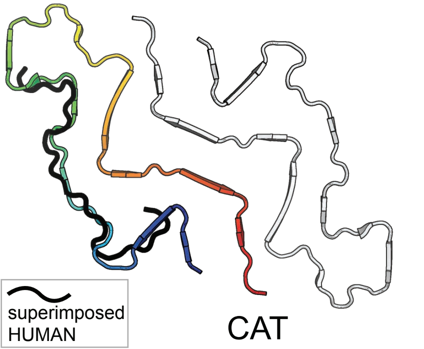

Figure S5

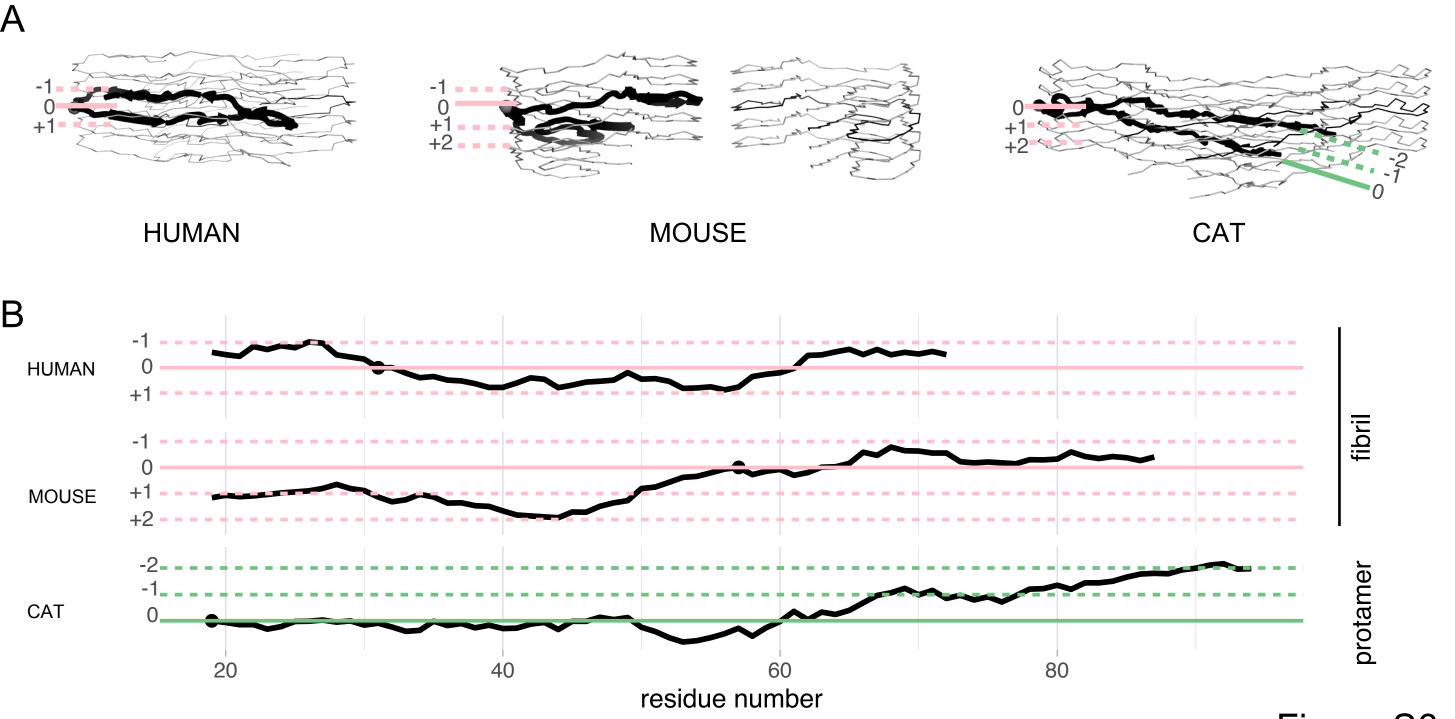
A
-1
0
+1
HUMAN
-1
0
+1
+2
MOUSE
0
+1
+2
-2
-1
0
CAT
B
HUMAN
-1
0
+1
MOUSE
-1
0
+1
+2
CAT
-2
-1
0
20
40
60
80
residue number
fibril
protamer

Figure S6

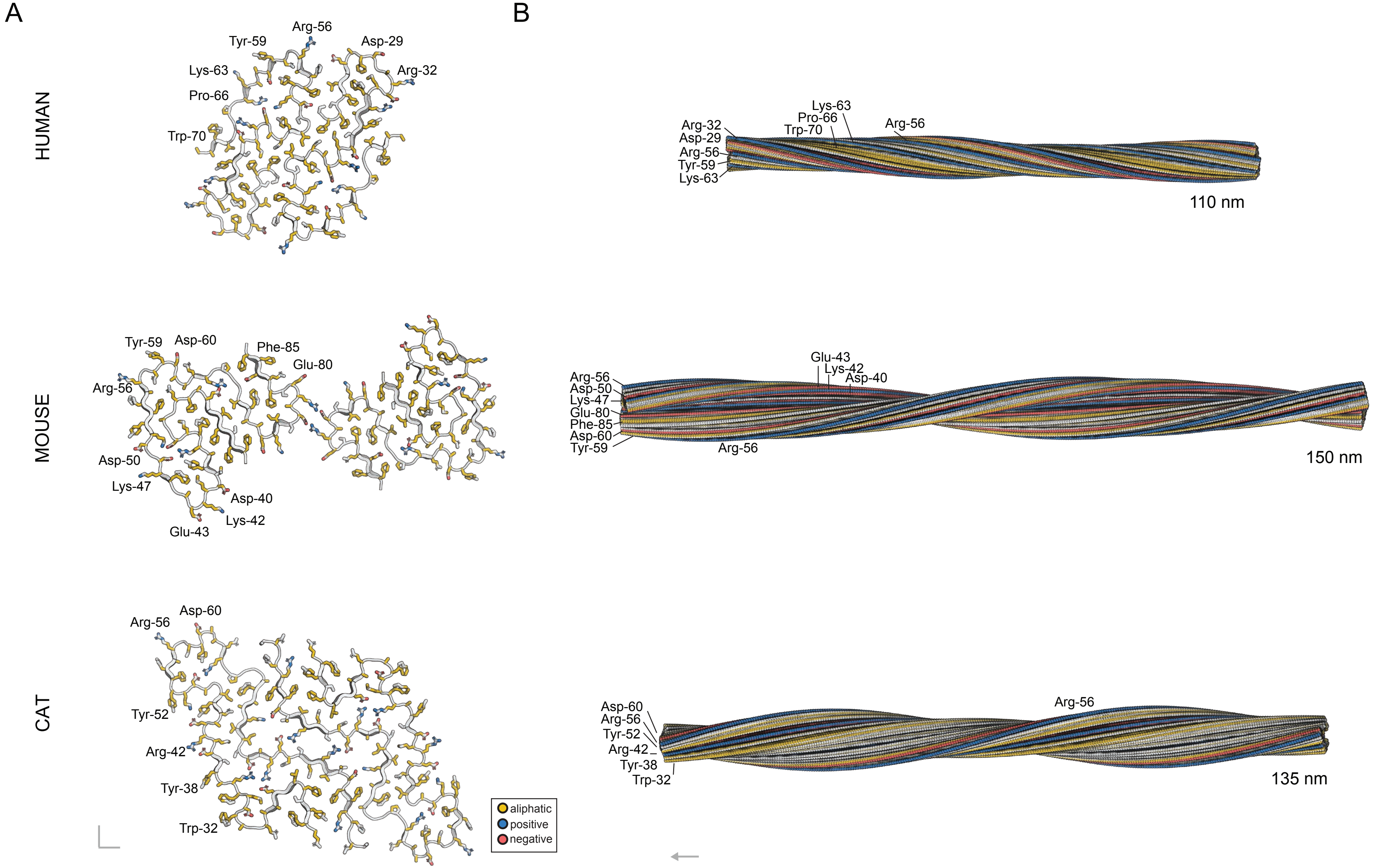

A
B
HUMAN
MOUSE
CAT
Arg-56
Tyr-59
Asp-29
Lys-63
Arg-32
Pro-66
Trp-70
Tyr-59
Asp-60
Phe-85
Glu-80
Arg-56
Asp-50
Lys-47
Asp-40
Lys-42
Glu-43
Arg-56
Asp-60
Tyr-52
Arg-42
Tyr-38
Trp-32
aliphatic
positive
negative
Lys-63
Pro-66
Arg-32
Trp-70
Arg-56
Asp-29
Arg-56
Tyr-59
Lys-63
110 nm
Glu-43
Lys-42
Asp-40
Arg-56
Asp-50
Lys-47
Glu-80
Phe-85
Asp-60
Tyr-59
Arg-56
150 nm
Arg-56
Asp-60
Arg-56
Tyr-52
Arg-42
Tyr-38
Trp-32
135 nm

Figure S7

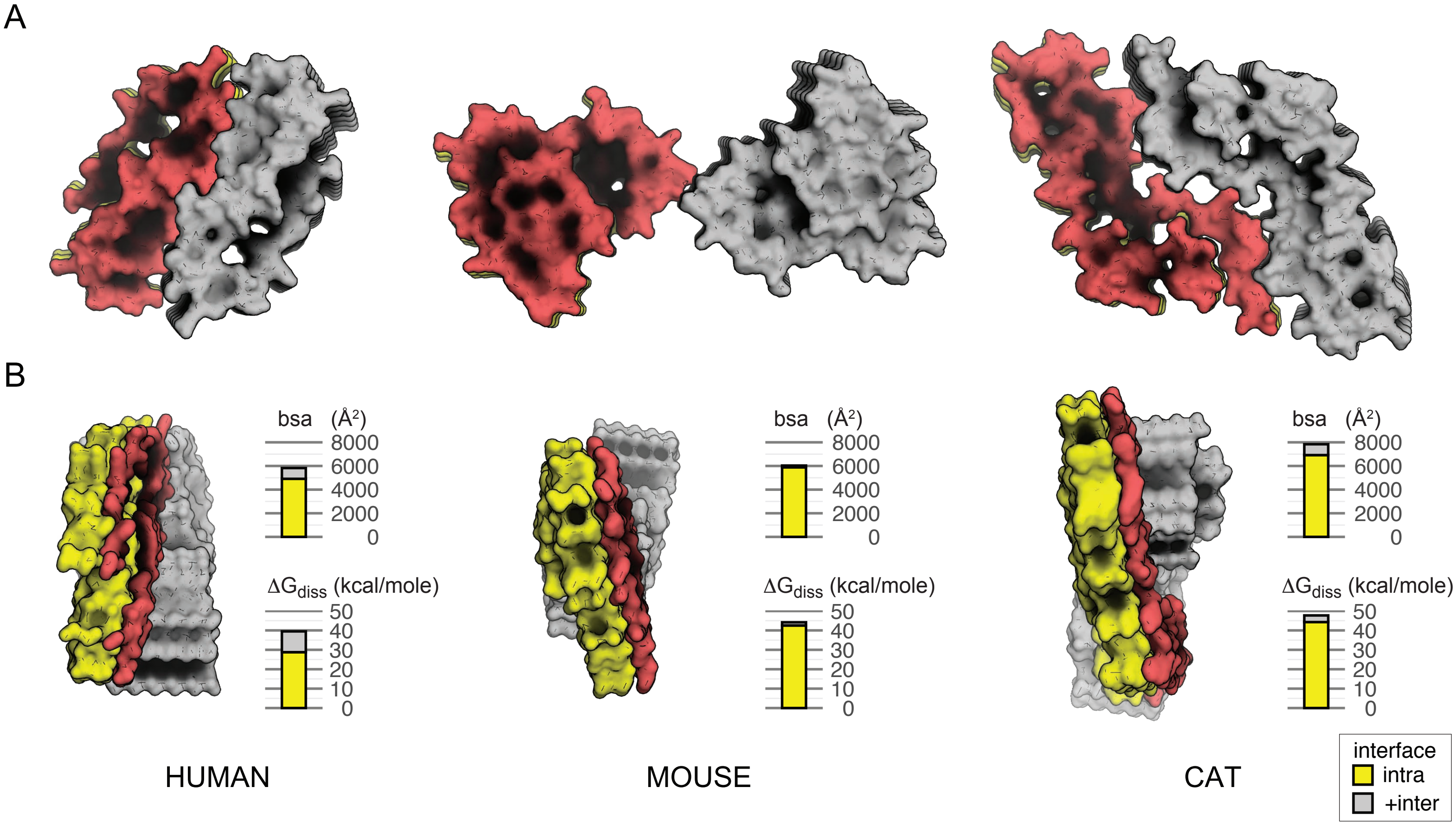
A
B
bsa (Å²)
8000
6000
4000
2000
0
ΔG_diss (kcal/mole)
50
40
30
20
10
0
HUMAN
MOUSE
CAT
interface
intra
+inter

Figure S8

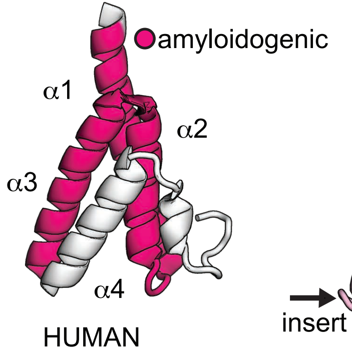

Figure S9
